# PIsToN: Evaluating Protein Binding Interfaces with Transformer Networks

**DOI:** 10.1101/2023.01.03.522623

**Authors:** Vitalii Stebliankin, Azam Shirali, Prabin Baral, Prem Chapagain, Giri Narasimhan

**Affiliations:** Bioinformatics Research Group (BioRG), Knight Foundation School of Computing and Information Sciences, Florida International University; 11200 SW 8th St, Miami, 33199, USA; Department of Physics, College of Arts, Science and Education, Florida International University; 11200 SW 8th St, Miami, 33199, USA; Biomolecular Sciences Institute, Florida International University; 11200 SW 8th St, Miami, 33199, USA

**Keywords:** Binding affinity, Protein complementarity, Protein interactions

## Abstract

The computational studies of protein binding are widely used to investigate fundamental biological processes and facilitate the development of modern drugs, vaccines, and therapeutics. Scoring functions aim to predict complexes that would be formed by the binding of two biomolecules and to assess and rank the strength of the binding at the interface. Despite past efforts, the accurate prediction and scoring of protein binding interfaces remain a challenge. The physics-based methods are computationally intensive and often have to trade accuracy for computational cost. The possible limitations of current machine learning (ML) methods are ineffective data representation, network architectures, and limited training data. Here, we propose a novel approach called PIsToN (evaluating **P**rotein binding **I**nterface**s** with **T**ransf**o**rmer **N**etworks) that aim to distinguish native-like protein complexes from decoys. Each protein interface is transformed into a collection of 2D images (interface maps), where each image corresponds to a geometric or biochemical property in which pixel intensity represents the feature values. Such a data representation provides atomic-level resolution of relevant protein characteristics. To build **hybrid** machine learning models, additional empirical-based energy terms are computed and provided as inputs to the neural network. The model is trained on thousands of native and computationally-predicted protein complexes that contain challenging examples. The multi-attention transformer network is also endowed with explainability by highlighting the specific features and binding sites that were the most important for the classification decision. The developed PIsToN model significantly outperforms existing state-of-the-art scoring functions on well-known datasets.

## 1 Introduction

With the advent of tools such as AlphaFold [29] that are based on machine learning, protein structure prediction has become more tractable. The next challenge in this domain is protein docking [7]. Given two protein structures (e.g., a designed molecule and a receptor or an antibody and an antigen), docking approaches aim to computationally predict the best binding location and conformation that could form a stable complex. Protein docking tools are critical for the successful development of drugs, vaccines, and therapeutics [64]. Virtual screening has been shown to save financial and labor resources in the drug design process [57]. However, modern computational tools for docking predict a large number of candidate complexes with good binding scores that fail confirmation in the laboratory [22].

The accuracy of molecular docking tools depends on the reliability of scoring functions. The first docking stage involves predicting many candidate conformations, either by fast Fourier transform (FFT) correlations or by direct search [23]. During the second stage, each docking conformation is ranked with a scoring function, and the top-ranked conformations are used as a final prediction [58]. The best docking tools have been able to ensure that the optimal binding conformation appears within the top 100 predicted docking structures, but rarely do they appear in the top 10, and even more rarely in the top position [45]. Furthermore, docking programs invariably predict binding sites for molecules that do not interact in nature, thereby generating a high false positive rate [1]. Therefore, improving scoring/evaluation functions would greatly benefit existing docking tools.

Accurate evaluation of protein complexes is a challenging problem [38]. A popular physics-based force field approach involves computing complex atomic interactions [33]. However, the physics-based method requires approximations due to the complexity of macro-molecular interactions, which result in reduced accuracy [41]. Another group of scoring functions called “empirical-based” include additional terms for better characterization of protein interactions, such as solvent accessible surface area (SASA) [16] and counts of the rotatable bonds [18]. The weights of the energy factors and protein characteristics are usually learned with linear regression [63]. Empirical-based scoring functions are generally better than force field methods but still yield poor correlations with experimental binding affinities and often fail to distinguish native interacting proteins from decoys [9].

Machine Learning (ML) provides promising approaches for improving the scoring accuracy of protein binding. An ML model can be viewed as a learnable function that maps protein complexes to their binding energies. First, unlike empirical-based methods, ML may learn nonlinear dependencies between energy terms. This is important given that noncovalent forces are interdependent in a nonlinear fashion [47]. Second, with good data embeddings, ML can escape the reliance on our incomplete knowledge of force fields and avoid propagating errors from approximations [40].

Several embeddings or representations for protein structures have been proposed for ML in the recent past. The first type includes using energy terms from empirical-based methods as a direct input to ML models, such as support vector machine [32, 12] or random forest [74]. Such models address non-linear dependency issues, but their accuracy remains limited by approximations from empirically-computed energy terms. The second set of methods includes features such as the frequency of each type of atom pairs that come within some threshold distance of each other [17, 4]. When combined with empirical-based energy terms, using contact frequency-based features is known to improve the scoring accuracy [37]. However, the results are still far from ideal, possibly because of the absence of 3D complementary information that can affect the stability of protein binding [39].

Other studies have incorporated detailed atomic spatial information by isolating a 3D voxel grid around the interface region and applied convolutional neural networks (CNNs) [68, 24, 3, 70, 56]. For example, DOVE constructs a voxel grid of atomic interaction types and their energetic contributions [70]. DeepRank used a 3D CNN architecture with extended atom- and residue-level features [56]. iScore uses innovative representations of protein interfaces through graph kernels [20]. While deep learning-based methods improve the accuracy of docking tools, they had a high rate of false positives among the top-ranked predictions [58]. In general, 3D CNNs are computationally expensive and prone to overfitting due to the sparsity of a voxel grid representation [34].

Several approaches have been proposed to reduce complex 3D information into compact signatures while preserving binding-related spatial features. For example, PatchBag characterized protein interface regions in terms of geometrical features from small surface units to search for evolutionary and functional relationships between proteins [6]. Deep Local Analysis evaluates the 3D conformational information with locally oriented cubes [46]. Molecular Surface Interaction Fingerprint (MaSIF) adapted a “patch” data representation to predict protein interactions [19]. A patch was defined as a region on a solvent-excluded protein surface with a fixed geodesic radius around a potential contact point. Each point on the surface is associated with geometric and physico-chemical features. A Siamese graph convolutional network was trained to minimize the distance between embeddings from interacting patches while maximizing it for non-binding ones. MaSIF reportedly is 1,000x faster than existing docking tools with only a slight loss of accuracy when compared to standard docking tools [19].

The limitations of MaSIF-Search include the following. First, the Siamese network can only take features belonging to a single protein as the input while excluding explicit interaction properties. While the Siamese approach allows ultra-fast scanning of molecular surface compatibility, it misses essential interaction terms, such as van der Waals forces, hydrogen bonds, desolvation, the distance between atoms of the opposite side, and many more. Second, the MaSIF training method generates negative patch pairs by randomly selecting surfaces outside the interface region. Therefore, negative instances consist of easily distinguishable non-complementary patches. We hypothesized that better training could be achieved using near-native non-interacting patch pairs. Third, the MaSIF network architecture consists only of convolutional layers, while better options such as attention-based models [72] and time series forecasting [59] could have been considered.

### Our contributions

Here we propose a tool called **P**rotein **I**nterface **S**coring with **T**ransf**o**rmer **N**etwork (PIsToN) that has the following novel contributions:

### Data representation

We represent interfaces of protein complexes as 2D multi-channel images. As in the MaSIF approach, circular “patches” on protein surfaces are first associated with geometric and physico-chemical features [19]. We perform an extra step of converting patches into images with pixel intensities corresponding to feature values associated with surface points at 1Å resolution. Unlike the single-patch approach of MaSIF, our method considers pairs of patches from protein binding interfaces, allowing us to compute essential interaction properties, such as the distance between atoms, relative accessible surface area (*RASA*), Van der Waals interactions, complementary surface charges and hydrophobicities, and more.

### Vision transformer network

We provide novel adaptations to the vision transformer (ViT) model, thus improving predictive performance and providing explainability, as explained below. Since ViTs are best suited for image classification [14], the choice of images for the representation of the features is an ideal complement.

### Multi-attention

In addition to the standard spatial attention of ViT, we appended another attention axis corresponding to a feature type (geometric or physico-chemical). The latent representations of each protein property were learned with an independent ViT network and combined in latent space using the transformer encoder.

### Hybrid machine learning

We enhanced the ViT model with a hybrid component that combines empirical energy terms with surface feature representations.

### Explainability

The multi-attention ViT allows for explainability in two ways: feature classes and binding sites essential for classification decision-making.

### Contrastive mini-batch training

We introduce a contrastive learning strategy with a novel loss function to learn discriminative embeddings for native binders and decoys. Prior approaches for scoring protein interfaces used either positive-random pairs [19] or a mixed batch of multiple positive and negative protein complexes [56, 20]. In our approach, each training iteration consists of multiple views of acceptable and incorrect binding poses of the same protein complex. The combination of supervised contrastive, margin ranking, and binary cross-entropy terms in the loss function help to cluster the correct docking models in the embedding space while pulling apart incorrect predictions.

### Superior Performance

As described later, the PIsToN model significantly outperforms state-of-the-art methods on two benchmark datasets.

## 2 Methods

### Interface maps

Given a protein complex structure in PDB format [5], we generate an *interface map* by projecting surface features from the binding interface of the two proteins to create a multi-channel image (Fig. 1), where each channel is dedicated to a feature type. Transforming data into 2D images (i.e., feature maps) prior to applying the ML model was successfully used for other ML applications by Valdes et al. [65]. Such transformations work well because ML techniques like convolutions work best on images.

**Fig. 1.**
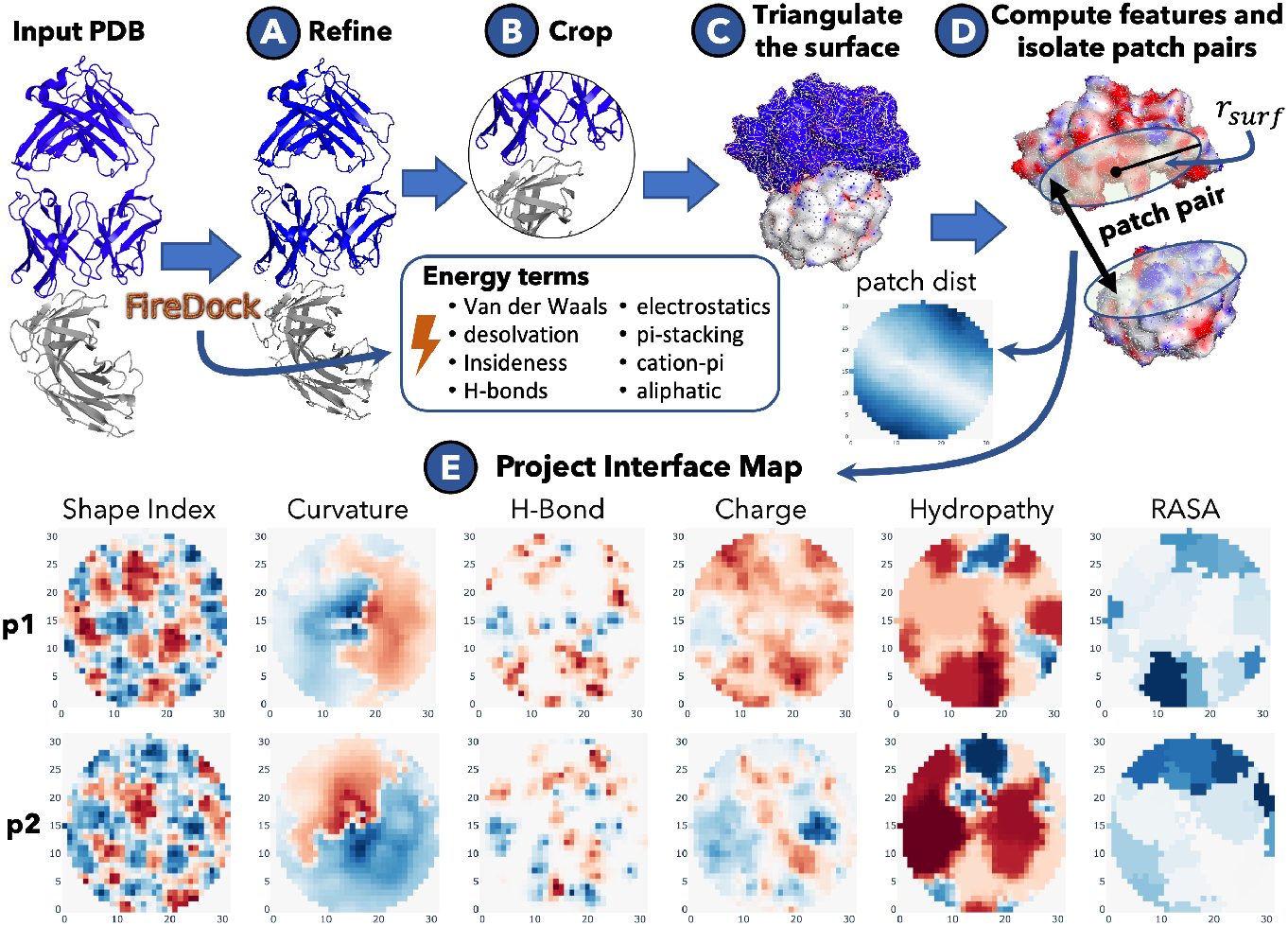
Feature engineering process for PIsToN (with an example on PDB ID 3B9K [60]). **A)**Executing the FireDock to refine a PDB structure and compute empirical energy terms. Cropping protein complexes by the radius *r*_*surf*_ near the protein interaction center. **C)** Triangulating the protein surface. **D)** Computing features associated with each point on a surface and isolating circular patches with the radius *r*_*surf*_. **E)** Projecting patch features into a 2D interface map.

Toward this goal, the protein structures were first refined to incorporate side-chain flexibility (Fig. 1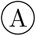). The restricted interface side-chain optimization with 50 Monte Carlo cycles was performed with FireDock (Fast Interaction Refinement in molecular Docking) [2]. The binding free energy terms of the refined structures computed by FireDock were set aside as an additional input to the deep learning model (see the center box in Fig. 1). The following energy terms were considered: van der Waals [21], desolvation [73], insideness [2], hydrogen and disulfide bonds [15], electrostatics [48], *π*-stacking, cation-*π*, and aliphatic interactions [11].

Second, the protein complexes were cropped to within distance *r*_surf_ from the “protein interaction center” (Fig. 1 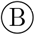), which was computed as the geometric center of contact points. Contact points were defined as the coordinates of atoms within 5Å proximity to an atom of the binding protein. Cropping improved memory efficiency by discarding residues with negligible influence on binding.

Third, the solvent-excluded surface was triangulated and re-scaled to a granularity of 1Å using *MaSIF* data preparation module [19] (Fig. 1 ©). Next, we computed features associated with a given pair of circular “patches” on the interacting surfaces (Fig 1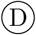). A patch is defined as the set of vertices on a triangulated protein surface within a geodesic distance of *r*_surf_ from the interaction center. We computed shape index, curvature, hydrogen bond potential, charge, and hydropathy features for each surface point on a surface using the MaSIF data preparation module [19]. Next, we computed another image for *patch dist*, which is generated from a grid of Euclidean distances between points on the two protein surfaces that project corresponding points on the patch pair. Lastly, the relative accessible surface area (*RASA*) [30] was computed for each patch residue using DSSP v2.3 [62].

Finally, patch features were converted into an image (interface map) with pixel intensities proportional to the feature values (Fig 1 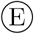). Surface points on patches were projected onto a 2D plane using multidimensional scaling (MDS) algorithm [43], which reduces the dimensionality to two while approximately preserving geodesic distances between two points on the patch. The final surface image of a given feature was computed as a 2D grid of size 2*r*_surf_ × 2*r*_surf_. The intensity of each pixel for a given feature *f* is equal to smoothed value over the four nearest neighbors:

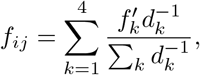

where *i, j* ∈ {0, 1, …, 2*r*_surf_} are the coordinates on the projected 2D grid, *f*_*ij*_ is the intensity of the pixel (*i, j*), *k* is the number of neighboring points on the protein surface close to (*i, j*), *d*_*k*_ is the Euclidean distance from (*i, j*) to the *k*-th surface point, and 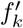 is the value of feature *f* of the *k*-th surface point.

### Network architecture

Fig. 2 shows ViT, Hybrid, and multi-attention components of the PIsToN network. The PIsToN*-ViT* was adapted from Vision Transformer architecture developed for image classification [14]. The input to the model was interface maps of size *a* projected from protein surfaces. Instead of conventional RGB, our ViT network accepts *N* channels corresponding to the number of surface features (Fig. 2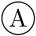). The image is subdivided into squares of size *l*, resulting in *P* = (*a/l*)^2^ mini-patches. Then, the collection of squares gets linearly projected into a set of *M*-dimensional token embeddings. Similar to the original ViT, we appended a learnable class token to the set of embeddings. The output of the transformer encoder is an embedded vector *Y* = {*y*_1_, *y*_2_, …, *y*_*M*_} and spatial attention map that highlight the mini-patches essential for the classification decision.

**Fig. 2.**
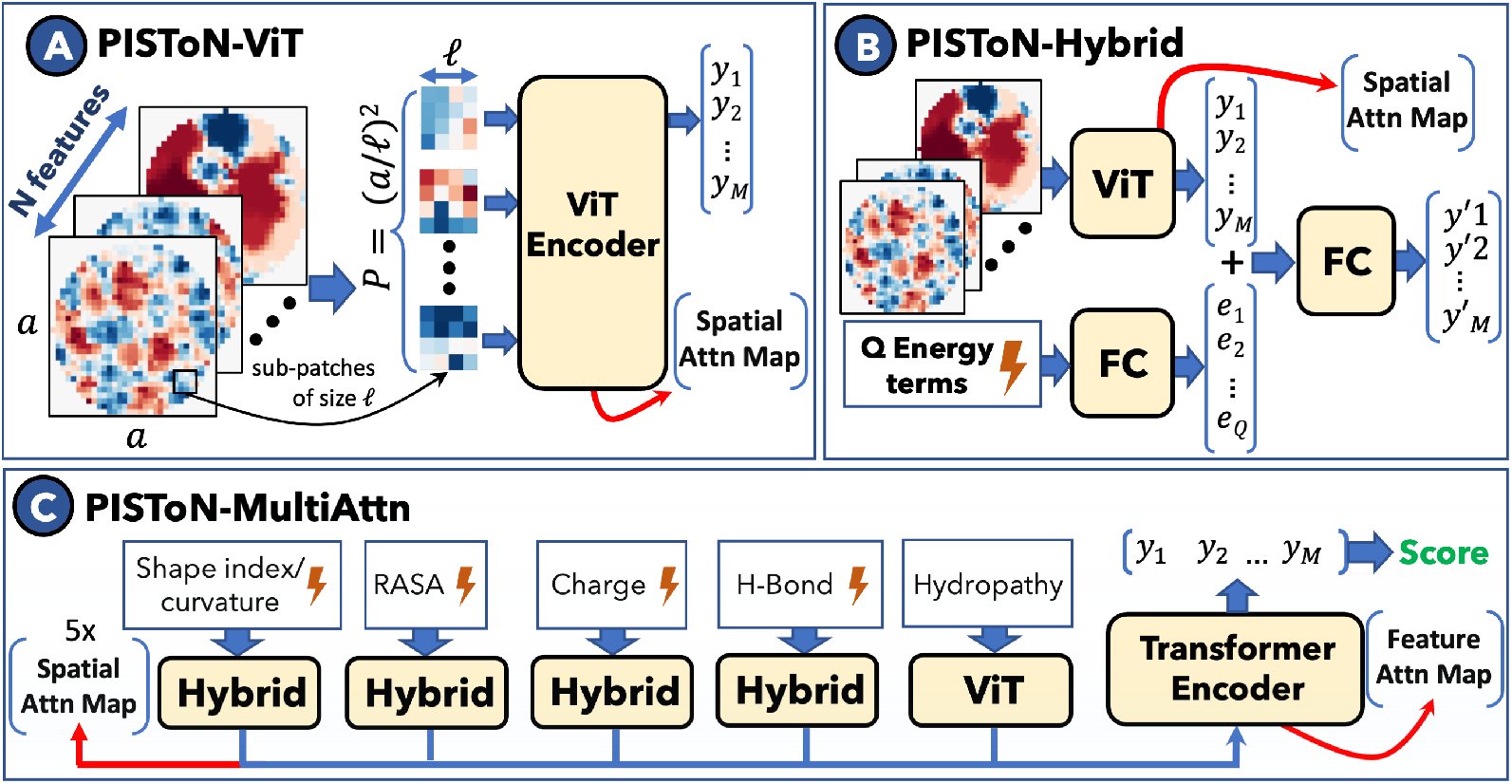
PIsToN Network architecture. **A)** ViT component of image classification to predict the binding class (“positive” or “incorrect”). **B)** Hybrid component that merges interface map with empirical energy terms. **C)** PIsToN Multi-attention network with parallel branches for each feature class that provides spatial and feature attention.

The PIsToN*-Hybrid* component was developed to combine empirically computed energies with the interface maps (Fig. 2 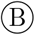). The *Q* energy terms were projected onto a latent vector {*e*_1_, *e*_2_, …, *e*_*Q*_} using a fully connected network (FC), which was then concatenated to the vector *Y* obtained from ViT. The merged vector was projected to a new vector *Y* ^*′*^ of size *M*, thus mapping the interface map and energy terms on to a latent space.

The PIsToN*-MultiAttn* efficiently combines the ViT and Hybrid components to include spatial and attention features. As shown in Fig. 2 ©, the input is subdivided into five branches of related interface maps and energy terms. Each branch contains an independent transformer network that learns separate latent vectors with spatial attention maps. The latent vectors of each branch are then aggregated with a class token into the transformer encoder for a final prediction. Also important is that feature attention map extracted to highlight the importance of each group. The first of the five feature groups is shape complementarity combined with Van der Waals (attractive and repulsive) and an “insideness” measure. Second, RASA was grouped with desolvation. Third, the charge feature was concatenated with electrostatic energies (attractive, repulsive, long range, and short range), pi-stacking, cation-pi, and aliphatic interactions. Fourth, H-bond interaction maps were combined with hydrogen and disulfide bonding energies. The last one was the hydropathy feature. Since the interatomic distances affect the energies from each group, the patch distance feature was appended as an extra channel to each branch. The class probability was computed as the Euclidean distance to the class centroid.

### Training process

For the training process, each native protein complex from the training set was separated and re-docked with HDOCK-light v. 1.1 to generate 100 docked models (Fig. S1 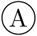) [71]. Each docked model was then compared with the native PDB structure to assess its quality. A docking model was labeled “acceptable” if it satisfied the following Critical Assessment of Predicted Interactions (CAPRI) condition [27]:

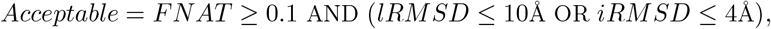

where *FNAT* is a fraction of native contacts recovered, *iRMSD* is the root mean square deviation (RMSD) between the interface *C*_*α*_ atoms of the docked model and the native structure, and *lRMSD* is the global RMSD between the *C*_*α*_ atoms of the two structures. Models for which the condition failed were labeled as “incorrect.” We believe that the way incorrect/negative examples are chosen is an important contributor to why our model performs better than MaSIF, which uses random patches to select the negative examples for training.

Training was performed in mini-batches, each consisting of multiple views of sets of acceptable and incorrect binding poses of a single protein complex (Fig. S1 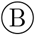). Since HDOCK produces many more incorrect docking models than acceptable ones, a mini-batch is limited to a 1:5 ratio of acceptable to incorrect models to limit the imbalance. Each epoch was trained on patches from the same *L* acceptable and 5*L* randomly selected incorrect docking poses. Each patch pair was augmented with random rotations from 0 to 360 degrees. The values of each pixel were normalized with standard scaling. The training was guided with a *contrastive learning* process (Fig. S1 ©). The loss function consisted of three terms – supervised contrastive [31], margin ranking [8], and binary cross entropy. The adaptations to the loss function are novel but due to space limitations, are explained in detail in the appendix.

The networks were trained with a maximum of 200 epochs with the early stopping condition of validation AUC not improving ten consecutive times. The *AdamW* optimizer with the learning rate of 0.0001 and weight decay of 0.001 was used for weights optimization [42]. The temperature *τ* that controls clustering tightness of contrastive loss was set to 0.5 [31].

### Datasets

The first dataset included the training and testing protein-protein interaction (PPI) complexes used by MaSIF software [19]. A minor fraction of complexes were excluded due to the memory overhead during the preprocessing step. In addition, 20% of the testing set was randomly subsampled into a validation set, which was used for the early stopping of PIsToN models. As a result, we preprocessed 4,942 training, 181 validation, and 678 testing native protein complexes. In total, we generated 17,291 positive and 452,031 negative training complexes. For validation, we obtained 703 positive and 16,864 negative docked structures. For testing, only one positive and one negative complex were randomly selected during the benchmark with other scoring functions, resulting in 678 positive and negative complexes. An additional 13 complexes from the CAPRI challenge set were used to test the PIsToN performance [36]. The protein docking models were extracted from the DeepRank study [56]. The dataset included 16,581 complexes generated by HADDOCK docking tool with corresponding CAPRI quality labels [13].

### Benchmarking

Our work was compared with a 2D CNN-based MaSIF-Search. The pre-trained MaSIF-Search “sc05” model was used with recommended parameters for data preprocessing [19]. Circular patches of 12Å radius were computed from the surfaces of interacting proteins using the MaSIF data preparation module. The data structure of the patch is a grid of 80 bins with five angular and 16 radial coordinates. Five features were extracted from each grid cell of a patch – shape index, distance-dependent curvature, electrostatics, hydropathy, and electrostatic potentials, resulting in an 80-dimensional descriptor vector. The network was trained such that the confidence of interaction between two patches is inversely proportional to the Euclidean distance between corresponding descriptors. The best minimum distance among the three most complementary patches in the complex was used as the measure for a binding score.

The performance on the CAPRI set was compared with three other deep learningbased methods: iScore [20], DOVE[70], and DeepRank [56]. The docking scores were extracted from the DeepRank study [56].

The empirical-based scoring functions can be subdivided into energy-based and potential-based methods. The binding score of the first category is defined as the weighted sum of energy terms. The following energy-based methods were considered: FireDock[2], PyDock[10], Rosetta[35], and ZRANK2[52]. Each scoring function includes electrostatics, desolvation, and van der Waals energy terms. FireDock and Rosetta include additional hydrogen and disulfide bonds. The unique term of FireDock is the internal energy that consists of bond stretching, angle bends, and torsional twists. The RosettaDock includes additional side chain rotamer energies. The potential-based methods involve computing the atomic and residue-level interaction properties, such as frequency of interaction types and solvent-accessible surface area. The following potential-based methods were considered: AP-PISA[67], SIPPER[53], and CP-PIE[54]. All empirical-based scoring functions we executed on CCharPPI online server[44].

### Evaluation metrics

The predictive power was estimated with the area under the receiver operating characteristic curve (AUC ROC) and the area under the Precision-Recall curve (AUPRC). The docking models with high, medium, and acceptable CAPRI quality were called “positive”, while the rest of the docking models were labeled “negative”. The ability to rank was evaluated with success rate, which is defined as the percentage of complexes for which at least one model of acceptable quality is found in the top *N* selected models. Other classification metrics included balanced accuracy, F1 score, precision, and recall. The classification threshold was chosen as the average of thresholds across 10-fold cross-validation with the maximum Matthews correlation coefficient, as proposed previously for the virtual screening application [1].

### Software and Hardware

The distance between atom coordinates was computed with KDTree from SciPy package v.1.1.0 [66]. The FNAT, iRMSD, and lRMSD were computed using pdb2sql Python v.0.5.2 [55]. The neural networks were implemented in PyTorch v.1.10 [51]. The interface maps were visualized with plotly v.5.10.0 [25]. The training and testing of deep learning models were performed on a GeForce GTX 1080 Ti 8 GPU, 256GB RAM, 28 core Intel Xeon CPU E5-2650 and also on IRCC resources at FIU [26].

## 3 Results

### Model selection

Each of the PIsToN components showed incremental improvement of AUC ROC based on the MaSIF test dataset (Table S1). The baseline AUC of the MaSIF architecture was 82.52% (Table S1, row 1). The CNN network trained with the concatenated 2D interface maps resulted in a competitive performance of AUC equal to 81.34% (Table S1, row 2). The CNN network had three blocks of convolutional and pooling layers, the sizes of which were constructed to match the MaSIF network (~66K trainable parameters). The vision transformer (ViT) with spatial attention (Fig. 2 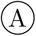) improved the AUC to 83.14% (Table S1, row 3). Inclusion of the additional features corresponding to Euclidean distances between points on the two protein surfaces from a patch pair (*patch dist*) and relative accessible surface area (RASA) improved the AUC to 87.85% (Table S1, rows 4-5). The inclusion of empirical energy terms via PIsToN hybrid component (Fig. 2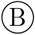) increased the AUC to 90.36% (Table S1, row 6). The multiattention architecture that matches energy functions with the corresponding physicochemical category of features (Fig. 2 ©) boosted the AUC by ~2.5% (Table S1, row 7).

The increase of the patch radius from 12Å to 16Å boosted the performance by another ~0.4% (Table S1, row 8). Finally, the implementation of the contrastive learning resulted in an AUC of ~93.55% (Table S1, row 9). Further increase of the patch radius from 16Å to 20Å did not improve the AUC, possibly due to overfitting (Table S1, row 10).

### Attention values and Explainability

The attention maps and the range of attention scores obtained from the PIsToN network allow us to analyze the essential features and binding sites that led to a classification decision. When applied to a MaSIF-test set, each feature branch of the PIsToN multi-attention component had a different distribution of attention values (Fig. 3, 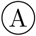). The network paid the greatest attention to geometric features (shape index and curvature) when predicting “positive” complexes (Fig. 3, 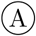, top) and to RASA when predicting complexes as decoys or “incorrect” (Fig. 3, 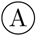, bottom).

**Fig. 3.**
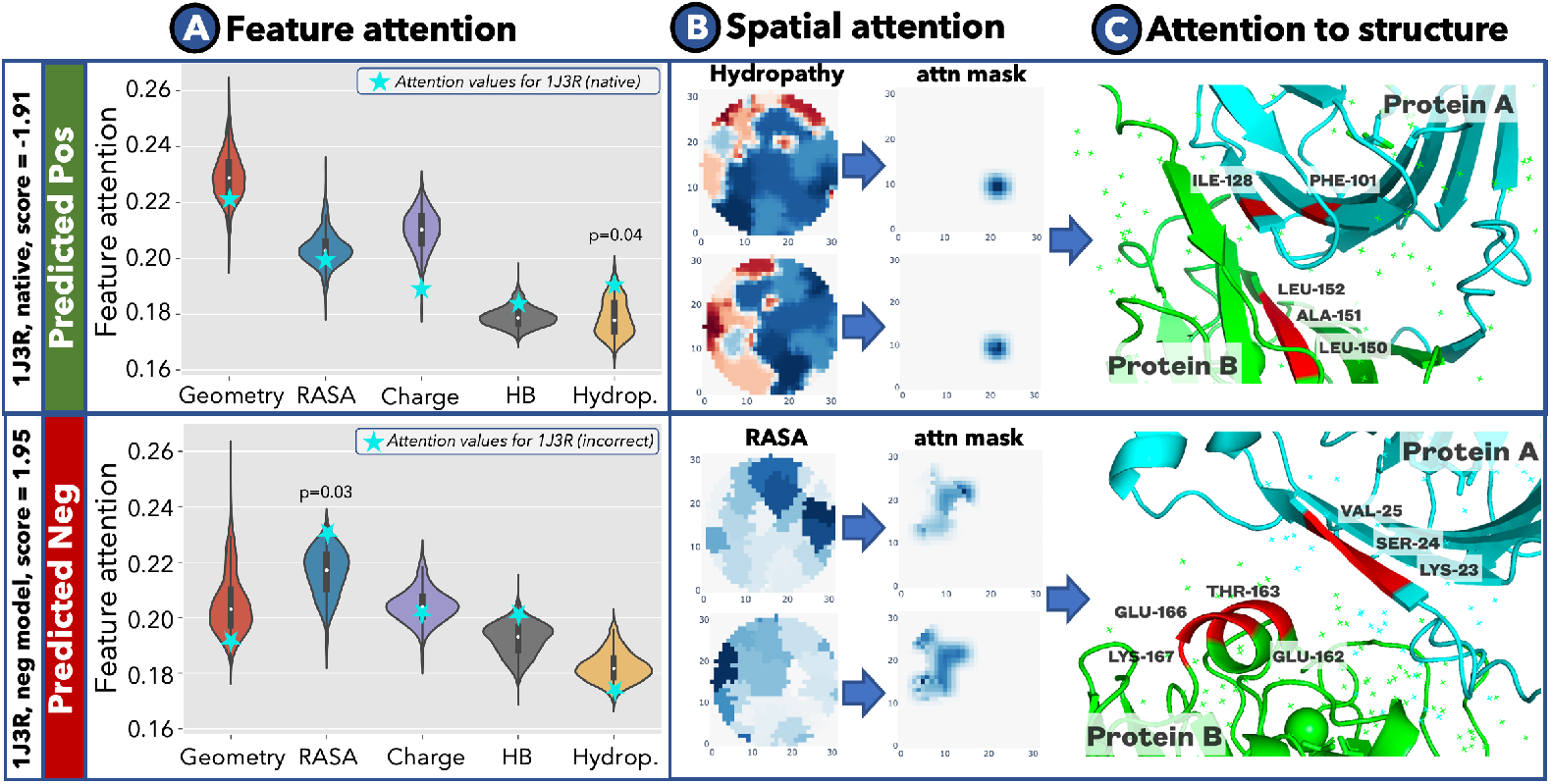
PIsToN explainability. **A)** Violin plots showing the distribution of attention values for five interface map features of complexes classified as positive (top) and incorrect (bottom). **B**An example of identifying significant pixels from spatial attention map for the 1J3R complex **C**Structure of 1J3R at the interface, where significant residues are highlighted in red. The feature attention values (A, turquoise stars), spatial attention maps (B), and structure (C) were shown for the 1J3R native complex (top) and incorrect docked model (bottom).

All attention values had similar numerical ranges (16–26%), suggesting the essentiality of each protein property for binding. However, the property could be of interest if the corresponding attention is in the upper tail of the distribution (Fig. 3, 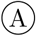). We considered a feature significant if the corresponding z-score of the attention value in the distribution is ≥ 1.96 (i.e., p-value ≤ 0.05). The same logic was applied to identify significant pixels from the spatial attention map.

Figure 3 shows an example of using attention maps for achieving explainability. The example is the homodimeric complex of the protein phosphogrucose isomerase (PDB ID 1J3R, chains A and B) from the archaeon *Thermococcus litoralis* [28]. Our network correctly predicted the native structure of the complex as “positive” with a score of −1.91.

The attention value for hydropathy was identified as significant (p-value = 0.04), while the rest of the features were within the normal distribution (Fig. 3, 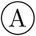, top, turquoise stars). We further explored the interface map of interacting proteins corresponding to hydropathy and identified the region of significant pixels from the spatial attention (Fig. 3, 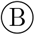, top). After mapping the pixels from the interface map to the original structure, we found that significant pixels corresponded to residues 128 (isoleucine) and 101 (phenylalanine) of chain A and 150-152 (leucine and alanine) of chain B (Fig. 3, ©, top). All of those residues are hydrophobic and located in the interaction region, suggesting a high role of the hydrophobic effect in the complex stability. Next, we analyzed a “decoy” docked model of the same protein pair (Fig. 3, bottom row). The PIsToN network predicted the score = 1.95, suggesting improbable binding. The network paid the most attention to a RASA feature (Fig. 3, 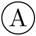, bottom row). The spatial attention module identified the larger region of significance (Fig. 3,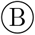, bottom row), which was close to the contact point on a complex (Fig. 3, ©, bottom row). The likely explanation is that the highlighted residues are surface accessible on both proteins but too far away, possibly destabilizing any complex that may be possible at that site.

### Comparison to Benchmark Tools

The PIsToN model showed a superior area under the receiver operating characteristic curve (AUC ROC), average precision (AP), balanced accuracy (BA), and F1-score when applied to MaSIF-test and CAPRI-score datasets (Table 1). The RoesettaDock [35] had the highest precision when applied to a MaSIF-test dataset and the lowest recall. Also, HADDOCK [13] had a higher recall on the CAPRI-score dataset but with low precision. Such abnormal precision and recall values could be the consequences of a suboptimal choice of a classification threshold.

**Table 1.** Classification performance of protein binding scoring functions. The results for the two datasets and the different methods included Area under the ROC (AUC ROC), average precision (AP), balanced accuracy (BA), F1-score, precision, and recall.

| Dataset | Method | AUC ROC | AP | BA | F1 | Precision | Recall |
| --- | --- | --- | --- | --- | --- | --- | --- |
| MaSIF-test | **PISToN** | **93.55** | **94.41** | **85.25** | **84.59** | 88.55 | **80.97** |
|  | MaSIF | 82.52 | 83.96 | 74.93 | 72.51 | 80.25 | 66.13 |
|  | FireDock | 89.95 | 90.54 | 81.83 | 81.58 | 82.81 | 80.38 |
|  | AP PISA | 89.85 | 90.30 | 81.98 | 81.28 | 84.63 | 78.18 |
|  | CP PIE | 88.11 | 89.28 | 81.03 | 79.94 | 84.87 | 75.55 |
|  | PyDOCK TOT | 86.11 | 87.17 | 77.66 | 77.62 | 77.79 | 77.45 |
|  | ZRANK 2 | 83.56 | 87.37 | 77.89 | 74.36 | 88.48 | 64.13 |
|  | RosettaDock | 75.28 | 82.77 | 76.06 | 69.81 | **94.50** | 55.34 |
|  | SIPPER | 74.52 | 72.23 | 68.64 | 70.32 | 66.80 | 74.23 |
| CAPRI-score | **PISToN** | **81.26** | **32.07** | **73.20** | **42.21** | **30.41** | 68.96 |
|  | iSCORE | 73.51 | 26.70 | 67.54 | 38.24 | 30.05 | 52.58 |
|  | DeepRank | 72.22 | 23.27 | 70.58 | 38.80 | 27.44 | 66.17 |
|  | HADDOCK | 70.35 | 28.67 | 69.88 | 36.63 | 24.77 | **70.31** |
|  | DOVE | 56.38 | 11.14 | 49.95 | 15.73 | 12.46 | 21.30 |

With PIsToN, we demonstrate a better chance of finding the correct model in the top *N* predictions. The success rate (percentage of complexes for which at least one model of acceptable quality is found in the top N selected models) of PIsToN in ranking the docking poses among many generated models (Table 2 and Fig. S4 E) is better. The benchmark contained 16,581 complexes generated by HADDOCK from 13 CAPRI targets, where at least two docked models are correct [56]. The PIsToN model were competitive with iScore to identify the correct docking pose as the top 1 prediction (Table 2). Starting from the top 10, PIsToN outperformed every other benchmarked scoring function.

**Table 2.**
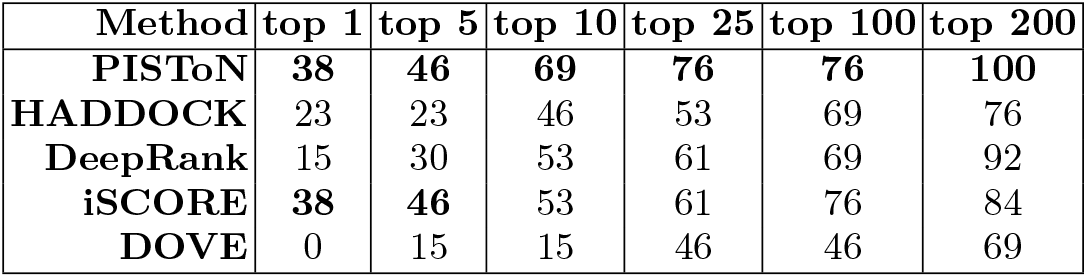
Success rate performance. The percentage of CAPRI targets for which at least one acceptable model is identified among top 1, top 5, top 10, top 25, top 100, and top 200 docking models.

| Method | top 1 | top 5 | top 10 | top 25 | top 100 | top 200 |
| --- | --- | --- | --- | --- | --- | --- |
| **PISToN** | **38** | **46** | **69** | **76** | **76** | **100** |
| **HADDOCK** | 23 | 23 | 46 | 53 | 69 | 76 |
| **DeepRank** | 15 | 30 | 53 | 61 | 69 | 92 |
| **iSCORE** | **38** | **46** | 53 | 61 | 76 | 84 |
| **DOVE** | 0 | 15 | 15 | 46 | 46 | 69 |

### Runtime

The PIsToN preprocessing and inferencing were more computationally efficient than MaSIF [19] (Fig. S3). PIsToN computes patches with a radius 12Å almost five times faster (Fig. S3,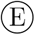, red and gray lines). Furthermore, PIsToN computed larger patches with the size of 16Å and 20Å faster than MaSIF’s 12Å patches (Fig. S3, 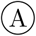, blue and purple lines). Such efficiency was achieved by removing (cropping) residues outside the interaction region that do not contribute to the properties of the computed patch. Similarly, PIsToN had a much faster inference time, possibly due to the memory efficiency of the PyTorch library (Fig. S3, 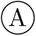). All runtime tests were performed on a single CPU and 256GB volume of RAM.

## 4 Discussion

The developed PIsToN deep learning model outperforms the current state-of-the-art empirical and machine learning-based protein scoring functions in identifying viable protein binding interfaces (Table 1). Our method introduces several novel elements. First, we represent molecular surface interfaces as machine learning-friendly multi-channel images, each corresponding to a geometric or biochemical property. (Fig. 1). Second, we introduce a novel hybrid multi-attention transformer network adapted from the popular image classification ViT network (Fig. 2) [14]. The hybrid component accepts empirically computed energy terms along with 2D interface maps. We adapted the ViT to include dual attention: spatial attention to highlight essential binding sites and feature attention to identifying the relative importance of specific protein properties (Fig. 3). Finally, we introduce a novel contrastive learning training strategy for protein interface classification. For each native protein structure, the network learns prototype centroids of acceptable and incorrect binding pose representations to achieve maximum separability between the classes (Fig. S1). Each innovative component showed incremental improvement in the model performance (Table S1).

The PIsToN outperforms other protein interface evaluation methods in classification and ranking tasks (Tables 1 and 2). However, the high rate of false positives for every method suggests the need for further improvements. We showed that the success rate for the top 1 prediction of scoring CAPRI models is as low as 38% for PIsToN and the other competitive tool, iScore [20] (Table 2). One possible reason for a limited ranking performance is the competition for binding sites. For example, if two proteins have several favorable binding spots, current approaches will fail to distinguish the one with the minimum energy. The PIsToN model predicts the viability of the protein binding interface and does not necessarily reflect the strength of the binding. The improvement can be achieved by additional training on the set of protein complexes with experimental binding affinities, such as PDBbind [69]. Yet, when the top 10 predictions were considered, the PIsToN identified 69% of correct complexes, which is significantly better than the competitors (Table 2).

The fact that PIsToN can rank the native binding complexes higher than other approaches suggests that our model will be of value in virtual screening. Given docking models for two protein targets, PIsToN has a higher chance of placing the correct configuration in the top 10 predictions, thus potentially speeding up screening of protein-protein interaction assays. PIsToN’s exceptional computational efficiency can save computational resources when thousands of candidates need to be screened (Fig. S3). While the current study focuses on evaluating macro-molecular binding, the PIs-ToN can be extended to protein-ligand interactions without any changes to the network. Another application can be molecular mimicry search, where a large set of antibody-antigen structures are scanned for cross-reactivity [61, 50].

The strong performance of PIsToN in comparison to other interface scoring methods suggests that it is not always necessary to invent new machine learning techniques for improved performance. However, it is far more important to use existing tools more effectively by engineering architectures that reflect our understanding of the domain.

## Acknowledgements

The authors thank the members of the Bioinformatics Research Group (BioRG) at FIU for their valuable feedback and comments. This work was supported by a grant from the National Science Foundation (CNS-2037374).

## Data availability

The lists of training/testing protein complexes and pre-computed interface maps are available at Zenodo (https://doi.org/10.5281/zenodo.7528053).

## Code availability

The PIsToN software and benchmark datasets are publicly available from https://biorg.cis.fiu.edu/piston/

## Appendix

**for “PIsToN: Evaluating Protein Binding Interfaces with Transformer Networks” by Stebliankin et al**.

### 1.1 Contrastive Learning

**Fig. S1.**
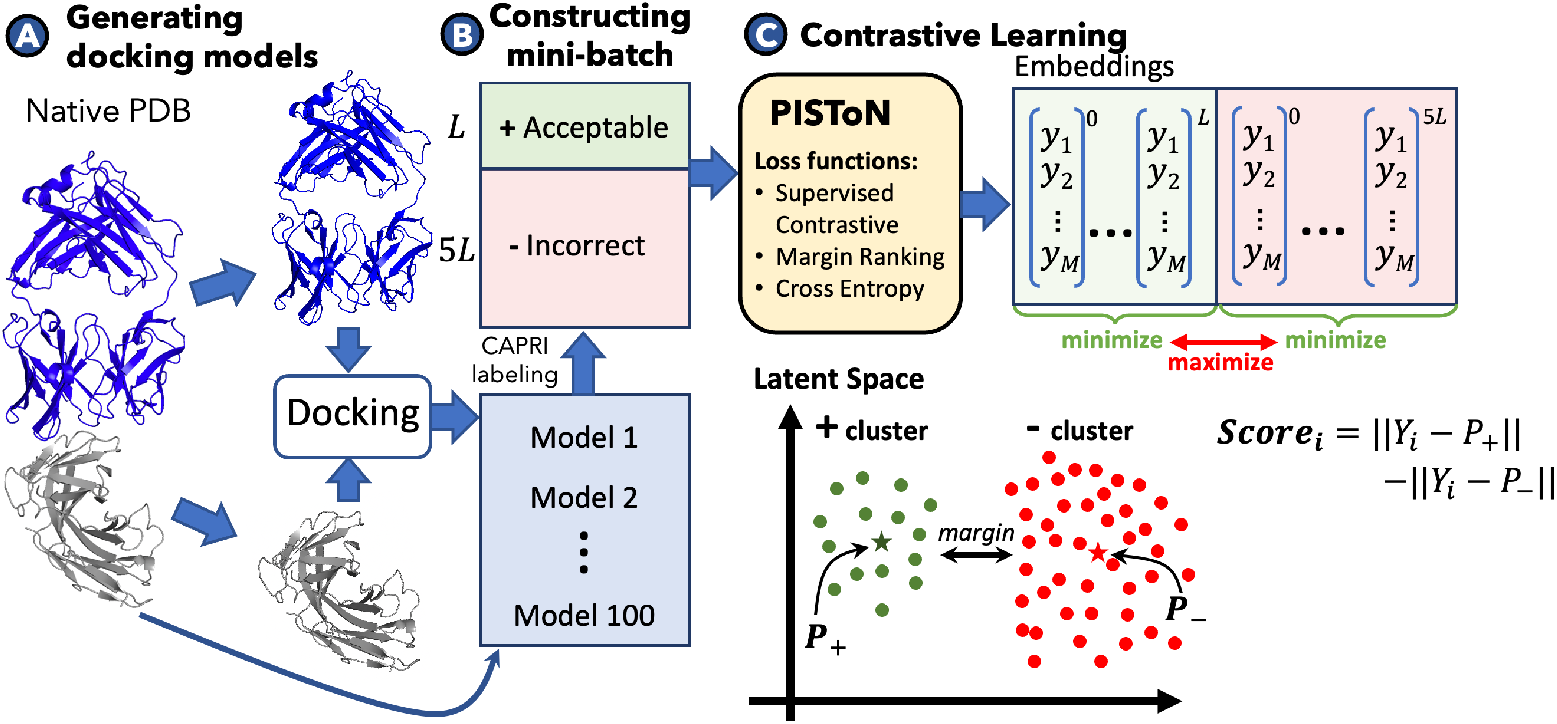
PIsToN training process. A) Generating 100 “acceptable” or “incorrect” docking models from a given native PDB structure from a training list. B) Constructing a mini-batch of “acceptable” and “incorrect” models with a 1:5 ratio; C) Update the weights of the PIsToN using contrastive loss functions to cluster embeddings of the same class near corresponding centroids while maximizing the distance between the clusters.

The training of PIsToN network was guided with a contrastive learning process (Supplementary Figure S1). The loss function consisted of three components: supervised contrastive (supCon), margin ranking, and binary cross entropy. Supervised contrastive loss was first introduced for image representational learning. The method aims to cluster instances of the same class in embedding space while simultaneously pushing apart clusters of samples from different classes [31]. The idea is similar to siamese training but generalized on an arbitrary number of positive and negative examples in a batch. In our formulations, the supervised contrastive loss looks as follows:

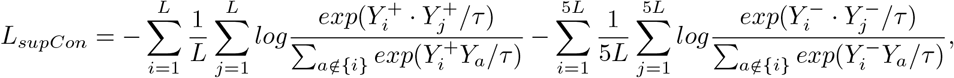

where L is the number of acceptable docking models, *Y*_*i*_ is the embedding for the *i*-th model of acceptable(+) or incorrect (−) protein complex, *a* is the set of instances excluding *i th* complex, and *τ* is the temperature hyperparameter that controls the clustering tightness.

To further coordinate separability, we introduced prototypical layers that serve cluster centroids (Figure S1 ©). The coordinates of positive and negative prototypes (*P* ^+^ and *P* ^−^) were learned with margin ranking loss:

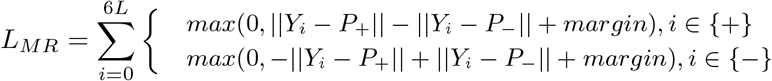

The interface score was predicted as the difference in distances to positive and negative prototypes, so that smaller values correspond to a better binding:

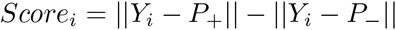

The final loss was the linear combination of contrastive, margin ranking, and binary cross entropy (BCE):

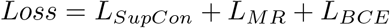

Since the loss function involves computing distance in high-dimensional feature space, we performed the L2 normalization of embedding vectors to transform the latent space into the unit hypersphere [49].

### 1.2 PIsToN scores

A PIsToN score for a given instance is computed as the difference in distances of the corresponding PIsToN embeddings of the positive and negative centroids (Fig. S1, C). Therefore, lower values correspond to the better binding. The maximum distance between two vectors in the latent space is two because the embeddings were normalized to a unit length (L2-normalization). Thus, the ideal binding corresponds to a score = −2 where the distance to a positive centroid is zero, and the distance to a negative centroid is two. In contrast, score=2 will correspond to the most unlikely binding interface.

The PIsToN scores accurately discriminated between native complexes and incorrect docking models (Fig. S2,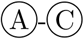). The high-quality docking predictions had a distribution of scores similar to native complexes (Fig. S2, 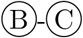, blue violins), while medium and acceptable categories had a larger overlap with incorrect models (Fig. S2, 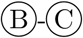, purple and gray violins). The distribution of scores for incorrect predictions in the CAPRI-score set was shifted to lower values (Fig. S2, ©, yellow violin), possibly due to the challenging nature of the dataset and a large number of incorrect models (Table S2).

The default classification threshold value for PIsToN is zero (Fig. S2, T1 dotted line) but can be adjusted depending on the dataset and the desired ratio between precision and sensitivity. To find the optimal classification boundary for MaSIF-test and CAPRI-score datasets, we computed the Matthews correlation coefficient (MCC) for different threshold values. The final classification threshold corresponded to the maximum MCC, averaged across ten splits, as suggested previously [1]. The threshold for the MaSIF-test dataset was 0.85, while a score of −0.5 was needed to best discriminate positive and negative complexes in the CAPRI-score dataset (Fig. S2,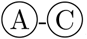, T1 and T3 dotted line). The same procedure for threshold optimization was performed for all other benchmark tools described in the next section (details in Table S3).

**Fig. S2.**
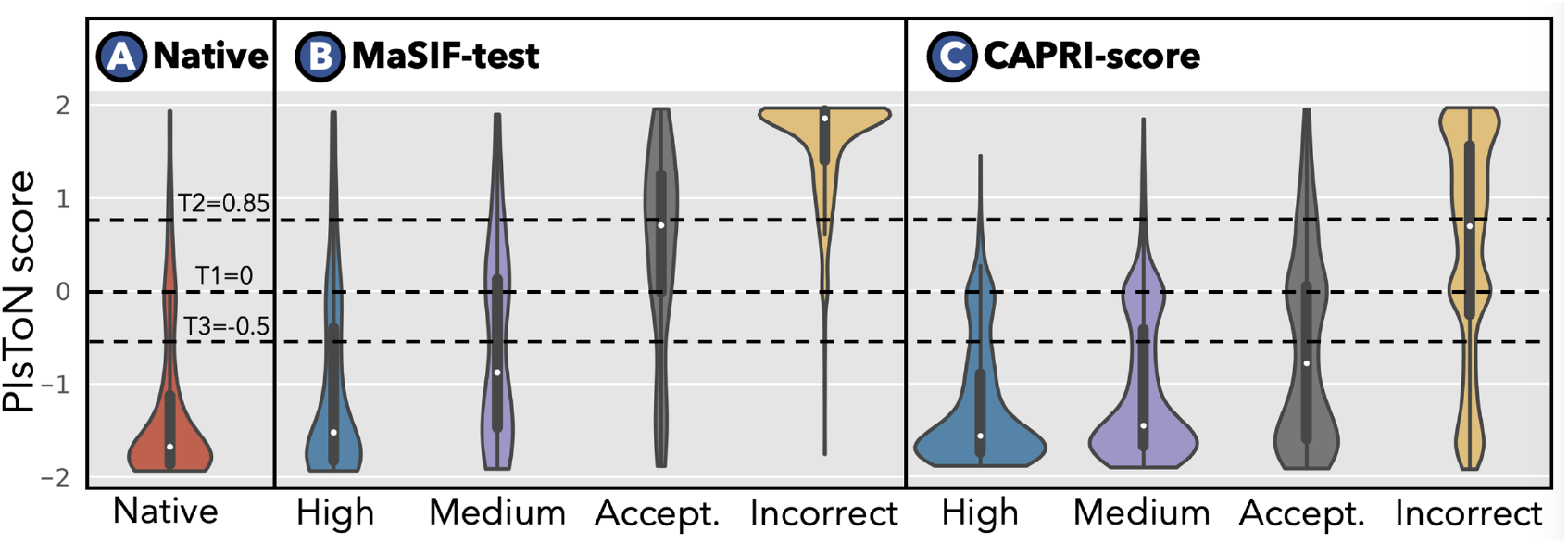
PIsToN **scores**. (**A**-**C**) Violin plots showing the distribution of PIsToN scores applied to three test sets: **A)** Native PDB complexes from the MaSIF-test set, **B)** docking models from MaSIF-test set, and **C)** Docking models from the CAPRI-score set. All docking models (B-C) are subdivided into “high”, “medium”, “acceptable”, and “incorrect” according to the CAPRI criteria. The dotted line shows our recommended thresholds: T1=0 if the dataset distribution is unknown, T2=0.85 if positive and negative instances are expected to be balanced, and T3=-0.5 if the data is expected to contain challenging examples with a high rate of false positives.

**Fig. S3.**
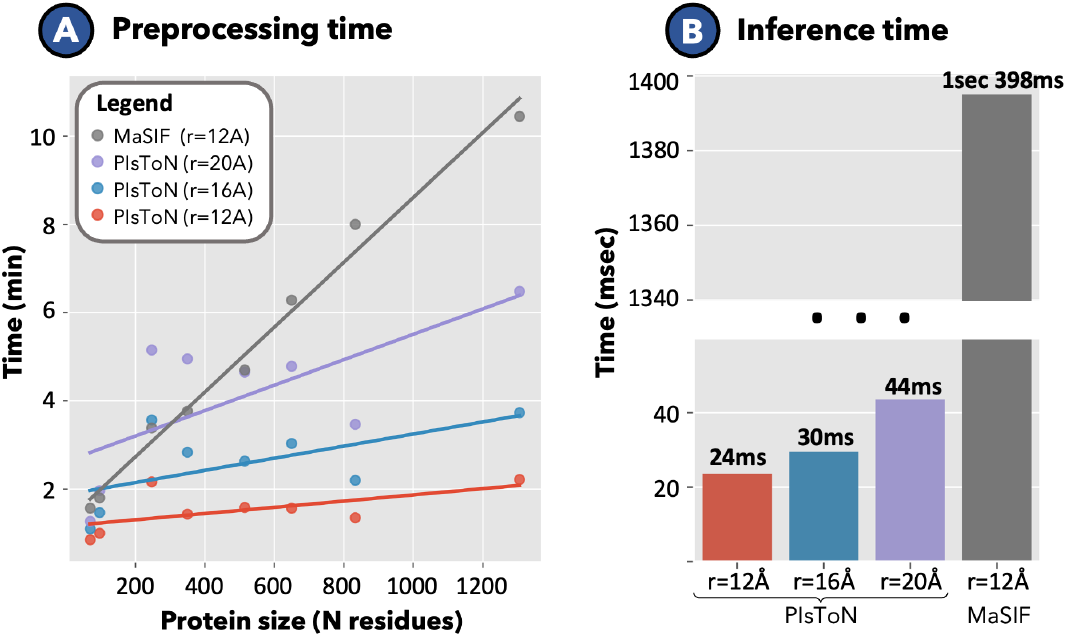
Runtime comparison of PIsToN and MaSIF. A) Prepossessing time to compute a patch pair for proteins of various sizes. B) Average inference time for a single protein.

**Fig. S4.**
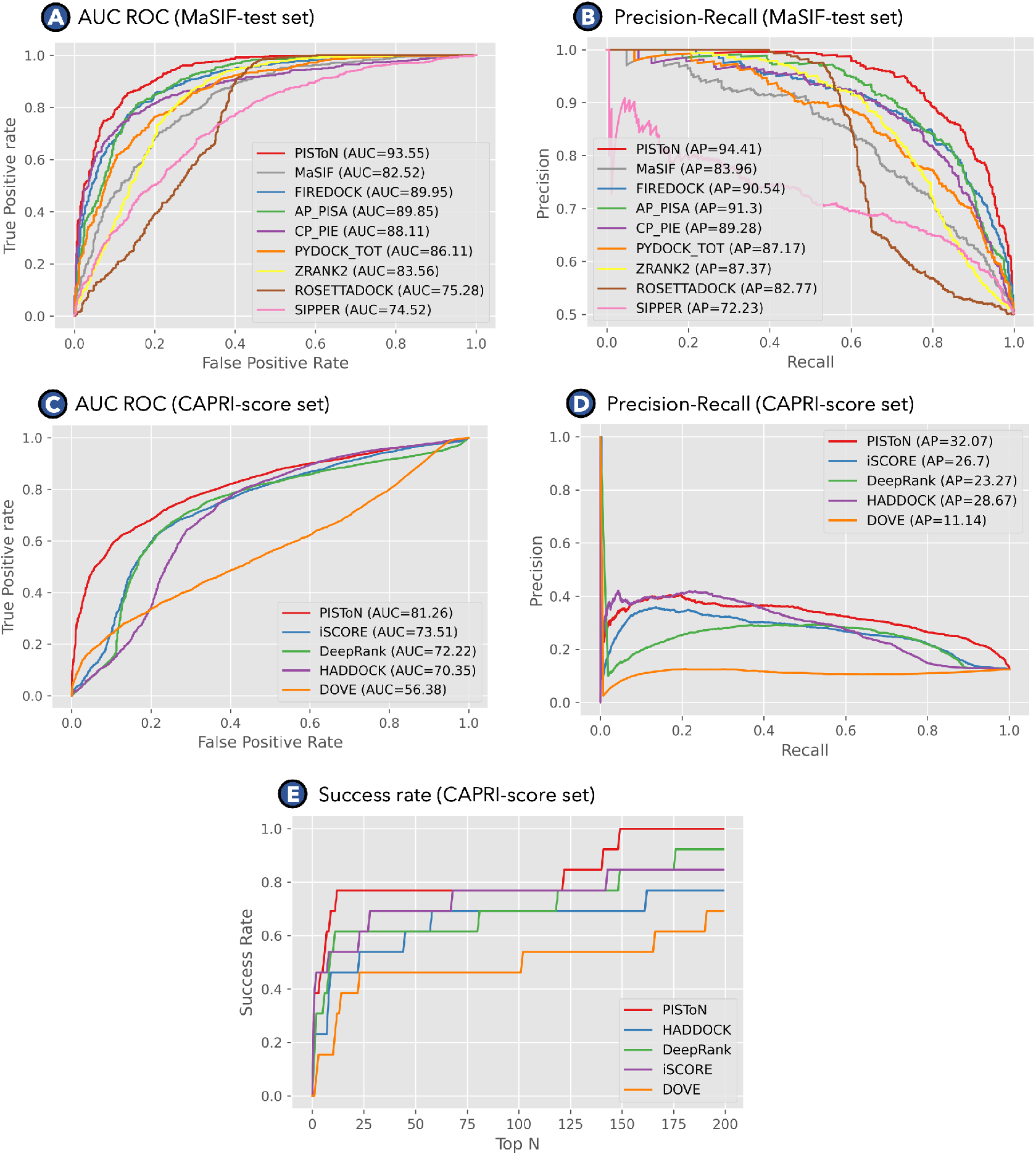
The performance of protein interface scoring functions A) ROC AUC on MaSIF test dataset; B) Precision-recall curve on MaSIF test dataset; C) ROC AUC on CAPRI score set; Precision-recall curve on CAPRI score set; E) The success rate on the CAPRI score set, computed as the percentage of complexes for which at least one model of acceptable quality is found in the top *N* selected models.

**Table S1.**
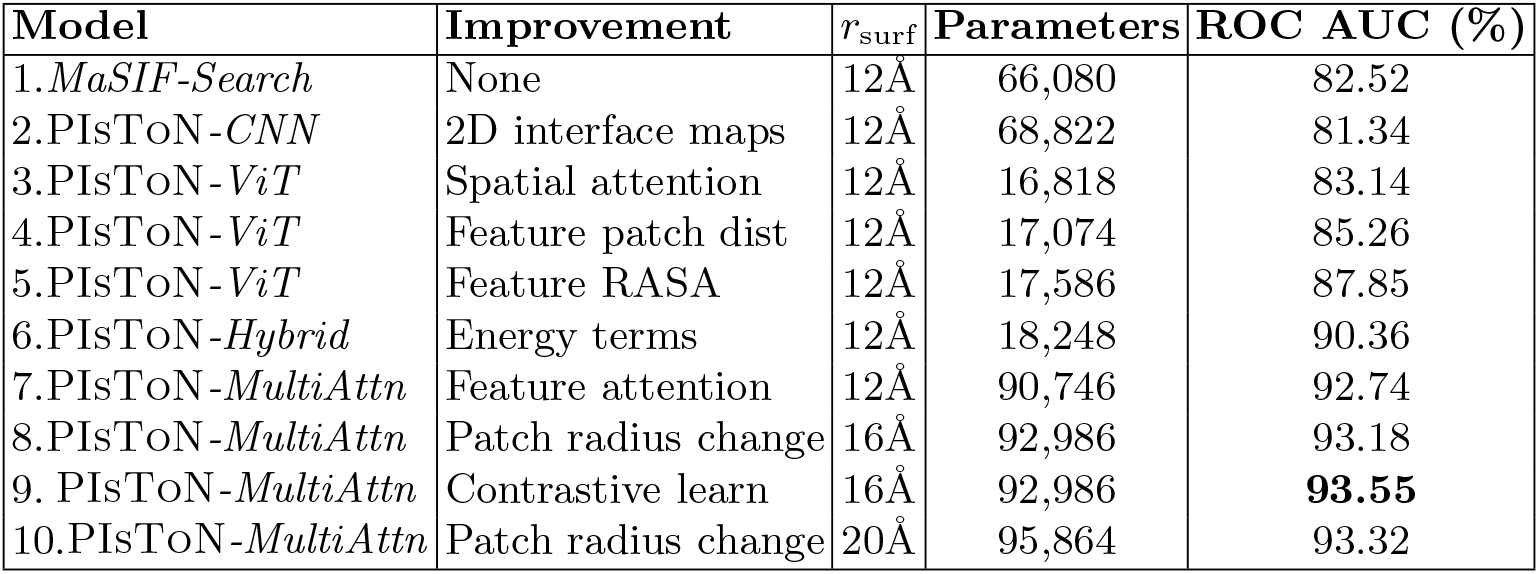
Performance improvement of PIsToN models evaluated on the MaSIF test dataset.

**Table S2.**
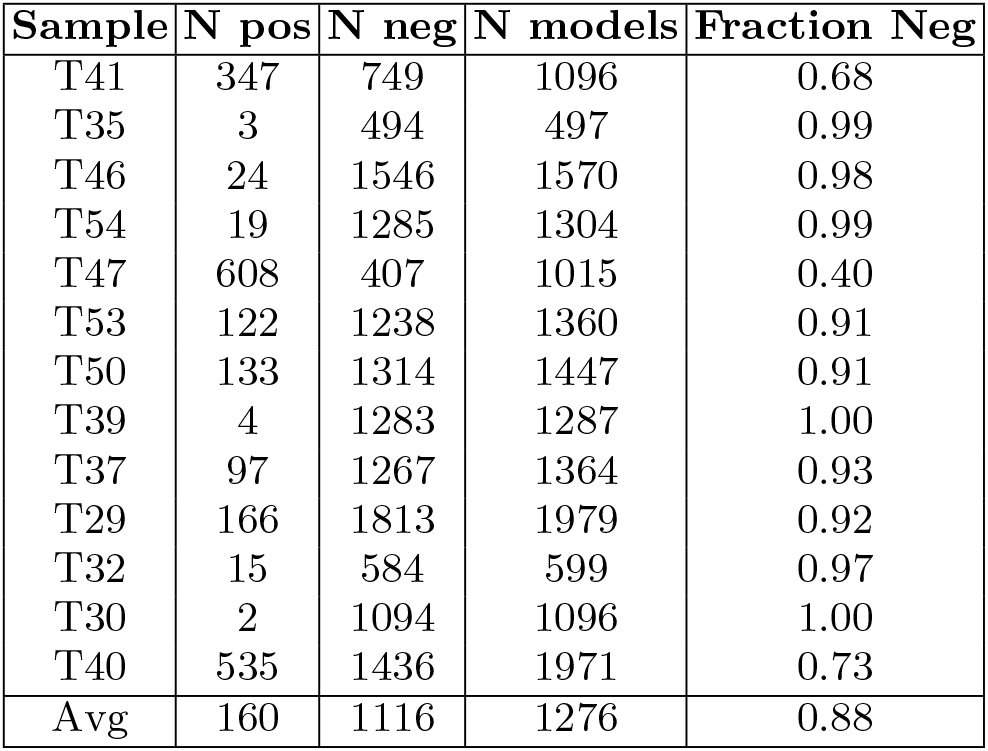
CAPRI-score dataset details. Each row shows the number of positive and negative complexes generated by HADDOCK for each CAPRI target.

**Table S3.**
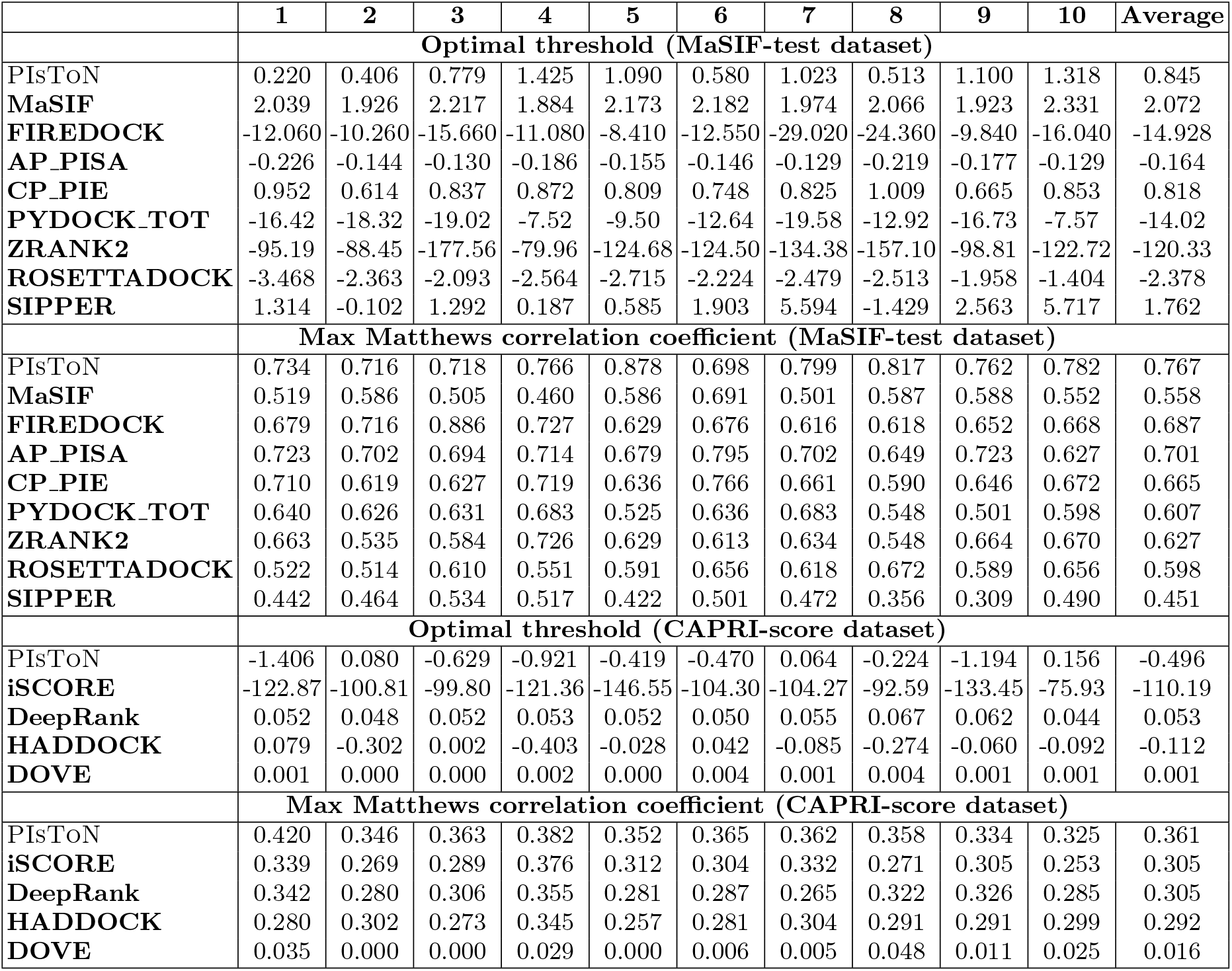
Finding an optimal threshold for scoring functions. Each optimal threshold corresponded to the maximum Matthews correlation coefficient averaged across 10-fold crossvalidation of the given testing set.

## Notes

https://biorg.cis.fiu.edu/piston/

https://zenodo.org/record/7528054

### Competing Interest Statement

The authors have declared no competing interest.

### Summary of Updates

We added a data availability section with referred Zenodo ID. A few sentences in the introduction, methods, and discussion were rephrased for clarification.

