## Supplementary materials for "PIsToN: Evaluating Protein Binding Interfaces with Transformer Networks"

### 1.1 Contrastive Learning

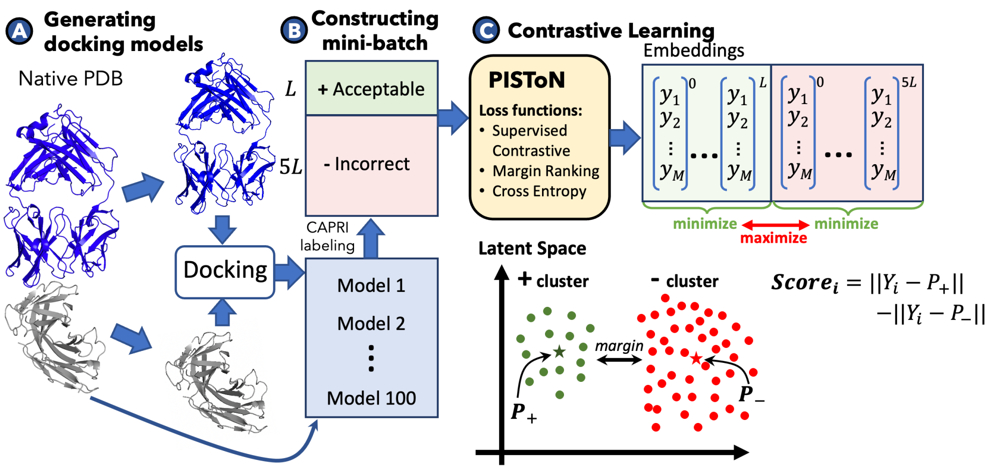


**Fig. S1.** PIsToN training process. A) Generating 100 "acceptable" or "incorrect" docking models from a given native PDB structure from a training list. B) Constructing a mini-batch of "acceptable" and "incorrect" models with a 1:5 ratio; C) Update the weights of the PIsToN using contrastive loss functions to cluster embeddings of the same class near corresponding centroids while maximizing the distance between the clusters.

The training of PIsToN network was guided with a contrastive learning process (Supplementary Figure S1). The loss function consisted of three components: supervised contrastive (supCon), margin ranking, and binary cross entropy. Supervised contrastive loss was first introduced for image representational learning. The method aims to cluster instances of the same class in embedding space while simultaneously pushing apart clusters of samples from different classes [31]. The idea is similar to siamese training but generalized on an arbitrary number of positive and negative examples in a batch. In our formulations, the supervised contrastive loss looks as follows:

$$L_{supCon} = -\sum_{i=1}^{L}\frac{1}{L}\sum_{j=1}^{L} log\frac{exp(Y_i^+ \cdot Y_j^+/\tau)}{\sum_{a\notin\{i\}} exp(Y_i^+ Y_a/\tau)} - \sum_{i=1}^{5L}\frac{1}{5L}\sum_{j=1}^{5L} log\frac{exp(Y_i^- \cdot Y_j^-/\tau)}{\sum_{a\notin\{i\}} exp(Y_i^- Y_a/\tau)},$$

where L is the number of acceptable docking models, $Y_i$ is the embedding for the $i$-th model of acceptable(+) or incorrect $(-)$ protein complex, $a$ is the set of instances excluding $i - th$ complex, and $\tau$ is the temperature hyperparameter that controls the clustering tightness.

To further coordinate separability, we introduced prototypical layers that serve cluster centroids (Figure S1 Ⓒ). The coordinates of positive and negative prototypes ($P^+$ and $P^-$) were learned with margin ranking loss:

$$L_{MR} = \sum_{i=0}^{6L} \begin{cases} max(0, ||Y_i - P_+|| - ||Y_i - P_-|| + margin), i \in \{+\} \\ max(0, -||Y_i - P_+|| + ||Y_i - P_-|| + margin), i \in \{-\} \end{cases}$$

The interface score was predicted as the difference in distances to positive and negative prototypes, so that smaller values correspond to a better binding:

$$Score_i = ||Y_i - P_+|| - ||Y_i - P_-||$$

The final loss was the linear combination of contrastive, margin ranking, and binary cross entropy (BCE):

$$Loss = L_{SupCon} + L_{MR} + L_{BCE}$$

Since the loss function involves computing distance in high-dimensional feature space, we performed the L2 normalization of embedding vectors to transform the latent space into the unit hypersphere [49].

#### 1.2 PIsToN scores

A PIsToN score for a given instance is computed as the difference in distances of the corresponding PIsToN embeddings of the positive and negative centroids (Fig. S1, Ⓒ). Therefore, lower values correspond to the better binding. The maximum distance between two vectors in the latent space is two because the embeddings were normalized to a unit length (L2-normalization). Thus, the ideal binding corresponds to a score $= -2$ where the distance to a positive centroid is zero, and the distance to a negative centroid is two. In contrast, score=2 will correspond to the most unlikely binding interface.

The PIsToN scores accurately discriminated between native complexes and incorrect docking models (Fig. S2, Ⓐ-Ⓒ). The high-quality docking predictions had a distribution of scores similar to native complexes (Fig. S2, Ⓑ-Ⓒ, blue violins), while medium and acceptable categories had a larger overlap with incorrect models (Fig. S2, Ⓑ-Ⓒ, purple and gray violins). The distribution of scores for incorrect predictions in the CAPRI-score set was shifted to lower values (Fig. S2, Ⓒ, yellow violin), possibly due to the challenging nature of the dataset and a large number of incorrect models (Table S2).

The default classification threshold value for PIsToN is zero (Fig. S2, T1 dotted line) but can be adjusted depending on the dataset and the desired ratio between precision and sensitivity. To find the optimal classification boundary for MaSIF-test and CAPRI-score datasets, we computed the Matthews correlation coefficient (MCC) for different threshold values. The final classification threshold corresponded to the maximum MCC, averaged across ten splits, as suggested previously [1]. The threshold for the MaSIF-test dataset was 0.85, while a score of $-0.5$ was needed to best discriminate positive and negative complexes in the CAPRI-score dataset (Fig. S2, Ⓐ-Ⓒ, T1 and T3 dotted line). The same procedure for threshold optimization was performed for all other benchmark tools described in the next section (details in Table S3).

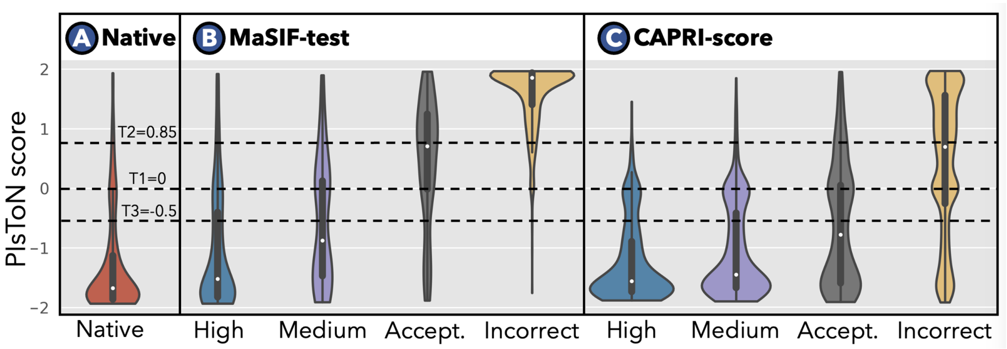


**Fig. S2.** PIsToN **scores.** (**A-C**) Violin plots showing the distribution of PIsToN scores applied to three test sets: **A)** Native PDB complexes from the MaSIF-test set, **B)** docking models from MaSIF-test set, and **C)** Docking models from the CAPRI-score set. All docking models (B-C) are subdivided into "high", "medium", "acceptable", and "incorrect" according to the CAPRI criteria. The dotted line shows our recommended thresholds: T1=0 if the dataset distribution is unknown, T2=0.85 if positive and negative instances are expected to be balanced, and T3=-0.5 if the data is expected to contain challenging examples with a high rate of false positives.

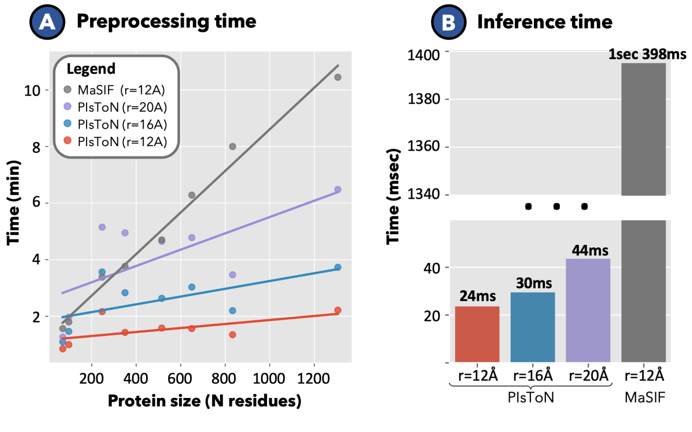


**Fig. S3. Runtime comparison of PIsToN and MaSIF.** A) Prepossessing time to compute a patch pair for proteins of various sizes. B) Average inference time for a single protein.

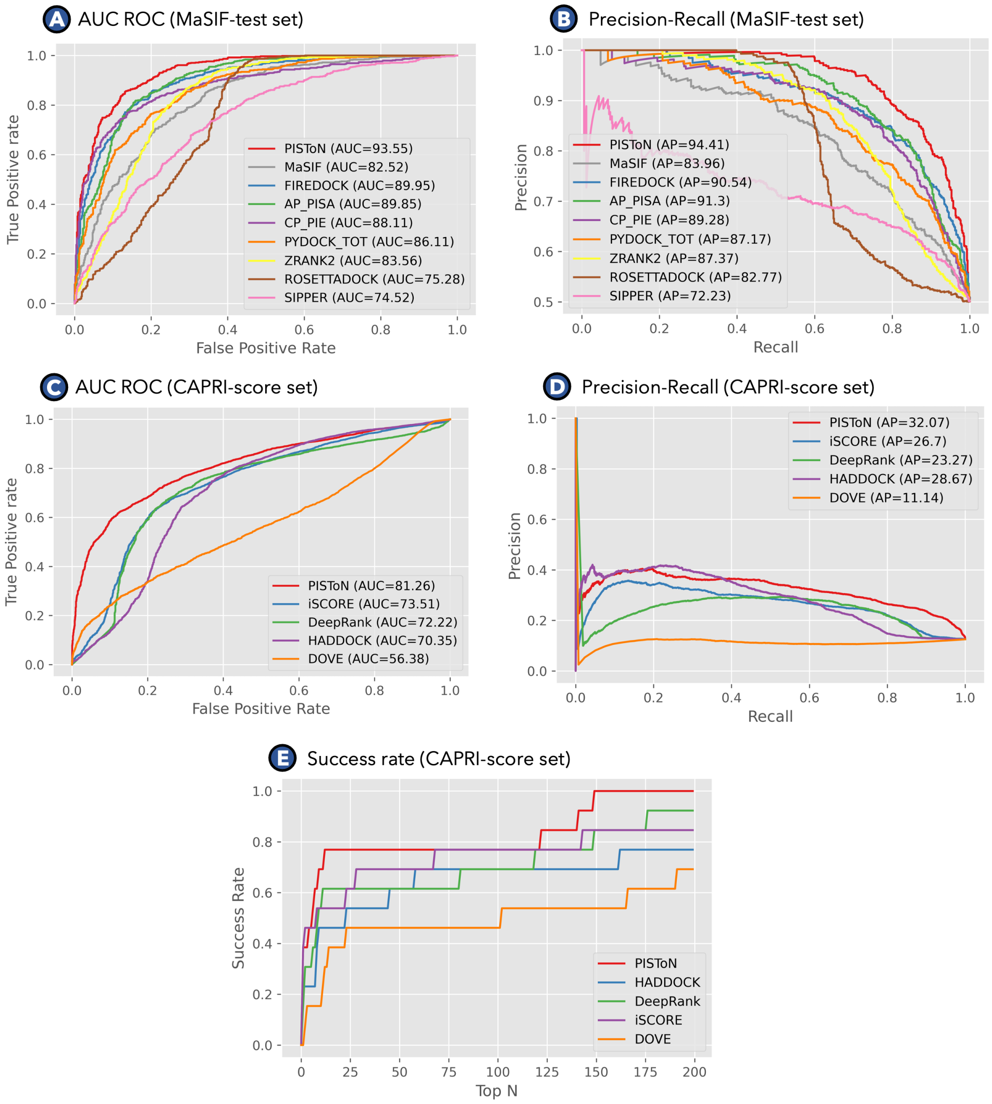


**Fig. S4.** The performance of protein interface scoring functions A) ROC AUC on MaSIF test dataset; B) Precision-recall curve on MaSIF test dataset; C) ROC AUC on CAPRI score set; D) Precision-recall curve on CAPRI score set; E) The success rate on the CAPRI score set, computed as the percentage of complexes for which at least one model of acceptable quality is found in the top $N$ selected models.

**Table S1.** Performance improvement of PIsToN models evaluated on the MaSIF test dataset.

| Model | Improvement | $r_{\text{surf}}$ | Parameters | ROC AUC (%) |
|---|---|---|---|---|
| 1.*MaSIF-Search* | None | 12Å | 66,080 | 82.52 |
| 2.PIsToN-*CNN* | 2D interface maps | 12Å | 68,822 | 81.34 |
| 3.PIsToN-*ViT* | Spatial attention | 12Å | 16,818 | 83.14 |
| 4.PIsToN-*ViT* | Feature patch dist | 12Å | 17,074 | 85.26 |
| 5.PIsToN-*ViT* | Feature RASA | 12Å | 17,586 | 87.85 |
| 6.PIsToN-*Hybrid* | Energy terms | 12Å | 18,248 | 90.36 |
| 7.PIsToN-*MultiAttn* | Feature attention | 12Å | 90,746 | 92.74 |
| 8.PIsToN-*MultiAttn* | Patch radius change | 16Å | 92,986 | 93.18 |
| 9. PIsToN-*MultiAttn* | Contrastive learn | 16Å | 92,986 | **93.55** |
| 10.PIsToN-*MultiAttn* | Patch radius change | 20Å | 95,864 | 93.32 |

**Table S2. CAPRI-score dataset details.** Each row shows the number of positive and negative complexes generated by HADDOCK for each CAPRI target.

| Sample | N pos | N neg | N models | Fraction Neg |
|---|---|---|---|---|
| T41 | 347 | 749 | 1096 | 0.68 |
| T35 | 3 | 494 | 497 | 0.99 |
| T46 | 24 | 1546 | 1570 | 0.98 |
| T54 | 19 | 1285 | 1304 | 0.99 |
| T47 | 608 | 407 | 1015 | 0.40 |
| T53 | 122 | 1238 | 1360 | 0.91 |
| T50 | 133 | 1314 | 1447 | 0.91 |
| T39 | 4 | 1283 | 1287 | 1.00 |
| T37 | 97 | 1267 | 1364 | 0.93 |
| T29 | 166 | 1813 | 1979 | 0.92 |
| T32 | 15 | 584 | 599 | 0.97 |
| T30 | 2 | 1094 | 1096 | 1.00 |
| T40 | 535 | 1436 | 1971 | 0.73 |
| Avg | 160 | 1116 | 1276 | 0.88 |

**Table S3. Finding an optimal threshold for scoring functions.** Each optimal threshold corresponded to the maximum Matthews correlation coefficient averaged across 10-fold cross-validation of the given testing set.

| | **1** | **2** | **3** | **4** | **5** | **6** | **7** | **8** | **9** | **10** | **Average** |
|---|---|---|---|---|---|---|---|---|---|---|---|
| | **Optimal threshold (MaSIF-test dataset)** | | | | | | | | | | |
| PIsToN | 0.220 | 0.406 | 0.779 | 1.425 | 1.090 | 0.580 | 1.023 | 0.513 | 1.100 | 1.318 | 0.845 |
| **MaSIF** | 2.039 | 1.926 | 2.217 | 1.884 | 2.173 | 2.182 | 1.974 | 2.066 | 1.923 | 2.331 | 2.072 |
| **FIREDOCK** | -12.060 | -10.260 | -15.660 | -11.080 | -8.410 | -12.550 | -29.020 | -24.360 | -9.840 | -16.040 | -14.928 |
| **AP_PISA** | -0.226 | -0.144 | -0.130 | -0.186 | -0.155 | -0.146 | -0.129 | -0.219 | -0.177 | -0.129 | -0.164 |
| **CP_PIE** | 0.952 | 0.614 | 0.837 | 0.872 | 0.809 | 0.748 | 0.825 | 1.009 | 0.665 | 0.853 | 0.818 |
| **PYDOCK_TOT** | -16.42 | -18.32 | -19.02 | -7.52 | -9.50 | -12.64 | -19.58 | -12.92 | -16.73 | -7.57 | -14.02 |
| **ZRANK2** | -95.19 | -88.45 | -177.56 | -79.96 | -124.68 | -124.50 | -134.38 | -157.10 | -98.81 | -122.72 | -120.33 |
| **ROSETTADOCK** | -3.468 | -2.363 | -2.093 | -2.564 | -2.715 | -2.224 | -2.479 | -2.513 | -1.958 | -1.404 | -2.378 |
| **SIPPER** | 1.314 | -0.102 | 1.292 | 0.187 | 0.585 | 1.903 | 5.594 | -1.429 | 2.563 | 5.717 | 1.762 |
| | **Max Matthews correlation coefficient (MaSIF-test dataset)** | | | | | | | | | | |
| PIsToN | 0.734 | 0.716 | 0.718 | 0.766 | 0.878 | 0.698 | 0.799 | 0.817 | 0.762 | 0.782 | 0.767 |
| **MaSIF** | 0.519 | 0.586 | 0.505 | 0.460 | 0.586 | 0.691 | 0.501 | 0.587 | 0.588 | 0.552 | 0.558 |
| **FIREDOCK** | 0.679 | 0.716 | 0.886 | 0.727 | 0.629 | 0.676 | 0.616 | 0.618 | 0.652 | 0.668 | 0.687 |
| **AP_PISA** | 0.723 | 0.702 | 0.694 | 0.714 | 0.679 | 0.795 | 0.702 | 0.649 | 0.723 | 0.627 | 0.701 |
| **CP_PIE** | 0.710 | 0.619 | 0.627 | 0.719 | 0.636 | 0.766 | 0.661 | 0.590 | 0.646 | 0.672 | 0.665 |
| **PYDOCK_TOT** | 0.640 | 0.626 | 0.631 | 0.683 | 0.525 | 0.636 | 0.683 | 0.548 | 0.501 | 0.598 | 0.607 |
| **ZRANK2** | 0.663 | 0.535 | 0.584 | 0.726 | 0.629 | 0.613 | 0.634 | 0.548 | 0.664 | 0.670 | 0.627 |
| **ROSETTADOCK** | 0.522 | 0.514 | 0.610 | 0.551 | 0.591 | 0.656 | 0.618 | 0.672 | 0.589 | 0.656 | 0.598 |
| **SIPPER** | 0.442 | 0.464 | 0.534 | 0.517 | 0.422 | 0.501 | 0.472 | 0.356 | 0.309 | 0.490 | 0.451 |
| | **Optimal threshold (CAPRI-score dataset)** | | | | | | | | | | |
| PIsToN | -1.406 | 0.080 | -0.629 | -0.921 | -0.419 | -0.470 | 0.064 | -0.224 | -1.194 | 0.156 | -0.496 |
| **iSCORE** | -122.87 | -100.81 | -99.80 | -121.36 | -146.55 | -104.30 | -104.27 | -92.59 | -133.45 | -75.93 | -110.19 |
| **DeepRank** | 0.052 | 0.048 | 0.052 | 0.053 | 0.052 | 0.050 | 0.055 | 0.067 | 0.062 | 0.044 | 0.053 |
| **HADDOCK** | 0.079 | -0.302 | 0.002 | -0.403 | -0.028 | 0.042 | -0.085 | -0.274 | -0.060 | -0.092 | -0.112 |
| **DOVE** | 0.001 | 0.000 | 0.000 | 0.002 | 0.000 | 0.004 | 0.001 | 0.004 | 0.001 | 0.001 | 0.001 |
| | **Max Matthews correlation coefficient (CAPRI-score dataset)** | | | | | | | | | | |
| PIsToN | 0.420 | 0.346 | 0.363 | 0.382 | 0.352 | 0.365 | 0.362 | 0.358 | 0.334 | 0.325 | 0.361 |
| **iSCORE** | 0.339 | 0.269 | 0.289 | 0.376 | 0.312 | 0.304 | 0.332 | 0.271 | 0.305 | 0.253 | 0.305 |
| **DeepRank** | 0.342 | 0.280 | 0.306 | 0.355 | 0.281 | 0.287 | 0.265 | 0.322 | 0.326 | 0.285 | 0.305 |
| **HADDOCK** | 0.280 | 0.302 | 0.273 | 0.345 | 0.257 | 0.281 | 0.304 | 0.291 | 0.291 | 0.299 | 0.292 |
| **DOVE** | 0.035 | 0.000 | 0.000 | 0.029 | 0.000 | 0.006 | 0.005 | 0.048 | 0.011 | 0.025 | 0.016 |
